# A novel dynamic network imaging analysis method reveals aging-related fragmentation of cortical networks in mouse

**DOI:** 10.1101/836817

**Authors:** Daniel A Llano, Chihua Ma, Umberto Di Fabrizio, Aynaz Taheri, Kevin A. Stebbings, Georgiy Yudintsev, Gang Xiao, Robert V. Kenyon, Tanya Y. Berger-Wolf

## Abstract

Network analysis of large-scale neuroimaging data has proven to be a particularly challenging computational problem. In this study, we adapt a novel analytical tool, known as the community dynamic inference method (CommDy), which was inspired by social network theory, for the study of brain imaging data from an aging mouse model. CommDy has been successfully used in other domains in biology; this report represents its first use in neuroscience. We used CommDy to investigate aging-related changes in network parameters in the auditory and motor cortices using flavoprotein autofluorescence imaging in brain slices and *in vivo*. Analysis of spontaneous activations in the auditory cortex of slices taken from young and aged animals demonstrated that cortical networks in aged brains were highly fragmented compared to networks observed in young animals. Specifically, the degree of connectivity of each activated node in the aged brains was significantly lower than those seen in the young brain, and multivariate analyses of all derived network metrics showed distinct clusters of these metrics in young vs. aged brains. CommDy network metrics were then used to build a random-forests classifier based on NMDA-receptor blockade data, which successfully recapitulated the aging findings, suggesting that the excitatory synaptic substructure of the auditory cortex may be altered during aging. A similar aging-related decline in network connectivity was also observed in spontaneous activity obtained from the awake motor cortex, suggesting that the findings in the auditory cortex are reflections of general mechanisms that occur during aging. Therefore, CommDy therefore provides a new dynamic network analytical tool to study the brain and provides links between network-level and synaptic-level dysfunction in the aging brain.

## Introduction

Normal aging is associated with a gradual loss of cognitive function [1–5]. The mechanisms responsible for this cognitive loss are not yet known, but given the increasing prevalence of aged individuals worldwide [6], it will be important to more fully understand the patterns of how brain networks fail with aging. Structural changes in the aging brain have been investigated and are characterized by changes in cortical thickness [7, 8], synaptic density [9–11] and selective loss of inhibitory interneurons [12, 13]. Less well characterized are functional changes in cortical physiology with aging, such as changes in functional connectivity. Functional connectivity between brain regions can change rapidly over time [14, 15], is not easily predictable from anatomical connectivity [16], and is altered in several different pathological states [17–19]. In addition, cortical networks appear particularly vulnerable to aging, and demonstrate diminished network-level functional connectivity over the lifespan [20, 21]. Furthermore, such aging-related disruptions in functional associations correlate with declines in cognitive performance [22]. However, current approaches to examine functional connectivity typically average data over long periods of time relative to timescales relevant for cognitive functions, and do not provide a clear mechanism to characterize the dynamics of functional connectivity over time. Therefore, techniques that are able to extract the dynamics of functional connectivity from brain imaging data will be of great value to those studying the impact of aging on the brain and to the broader neuroscience community.

Network analysis is an emerging approach to understand functional connectivity of the brain [23–25]. Unfortunately, network analysis of brain imaging data has proven to be a particularly challenging computational problem. Brain activity is intrinsically highly dynamic, whereby functional associations between neurons and brain regions ebb and flow as the organism’s level of arousal or focus of attention change. Since most previous network methodologies rely on collecting time series over long periods of time to build maps [26–28], the dynamic information extracted from these networks is lost. There have been recent attempts to ameliorate this problem by examining functional connectivity over time. These approaches have generally consisted of choosing a window of time for analysis and sliding this window along the period of data acquisition [29, 30]. Some of the problems with this approach have been: 1) the analysis windows (typically about 30-60 seconds) are still long relative to the cognitive processes of interest which evolve over hundreds of milliseconds, 2) choosing smaller windows tends to diminish signal-to-noise ratios and 3) noise may be non-stationary and therefore introduce spurious “networks” along the time course of analysis [14].

Previous investigators have suggested that complex dynamic networks, be they populations of neurons, large ecosystems, or social networks, share common underlying organizational motifs [23, 31–33]. Therefore, common methods may be used to describe and understand these networks. One dynamic network analysis method, known as the community dynamic inference method (CommDy) [34], which is grounded in social network theory, groups nodes into communities (functional clusters) in a way that minimizes the overall changes in the community affiliations of individuals over time. CommDy was developed specifically to characterize dynamics and was originally applied to the study of social networks, which is an area of study that has been plagued by the same conceptual and computational bottlenecks seen in brain imaging data analysis. For example, CommDy has successfully characterized group behavior of sheep [35] and equids [36], as well as social interactions among groups of people [37], though it has not yet been applied to neuroscience.

In the current report, we adapt CommDy for the analysis of flavoprotein autofluorescence imaging data from the mouse cerebral cortex in young and aged animals obtained using both brain slice and whole-animal approaches. Flavoprotein autofluorescence imaging capitalizes on intrinsic fluorescence that occurs in mitochondrial flavoproteins as neurons become active. We and others have found that such signals are very stable over time and are highly sensitive to neuronal activation [38–40]. Here, we examine changes in the auditory and motor cortices. These brain regions were chosen based on ease of data acquisition, but have both been shown to show significant vulnerability with aging [41–46]. Despite our growing understanding of the changes that occur in the cortex with aging, we lack a network-level description of how aging impacts the cerebral cortex and what synaptic changes may give rise to these network-level differences. Therefore, in this study, we adapt CommDy for the analysis of neuroimaging data and compare network-level changes in the auditory and motor cortices of young and aged mice.

## Results

### Method development

Two datasets were examined in this study: brain slice imaging data from the auditory cortex during paroxysmal depolarizations and *in vivo* imaging data from the awake mouse motor cortex (See Figure 1). These regions and types of preparations were used for reasons of experimental convenience and illustrate the flexibility of the CommDy technique to handle different types of datasets. Initial development of CommDy was done on slice data, where the signal-to-noise ratios are high, and then adapted for *in vivo* spontaneous activity data.

**Figure 1:**
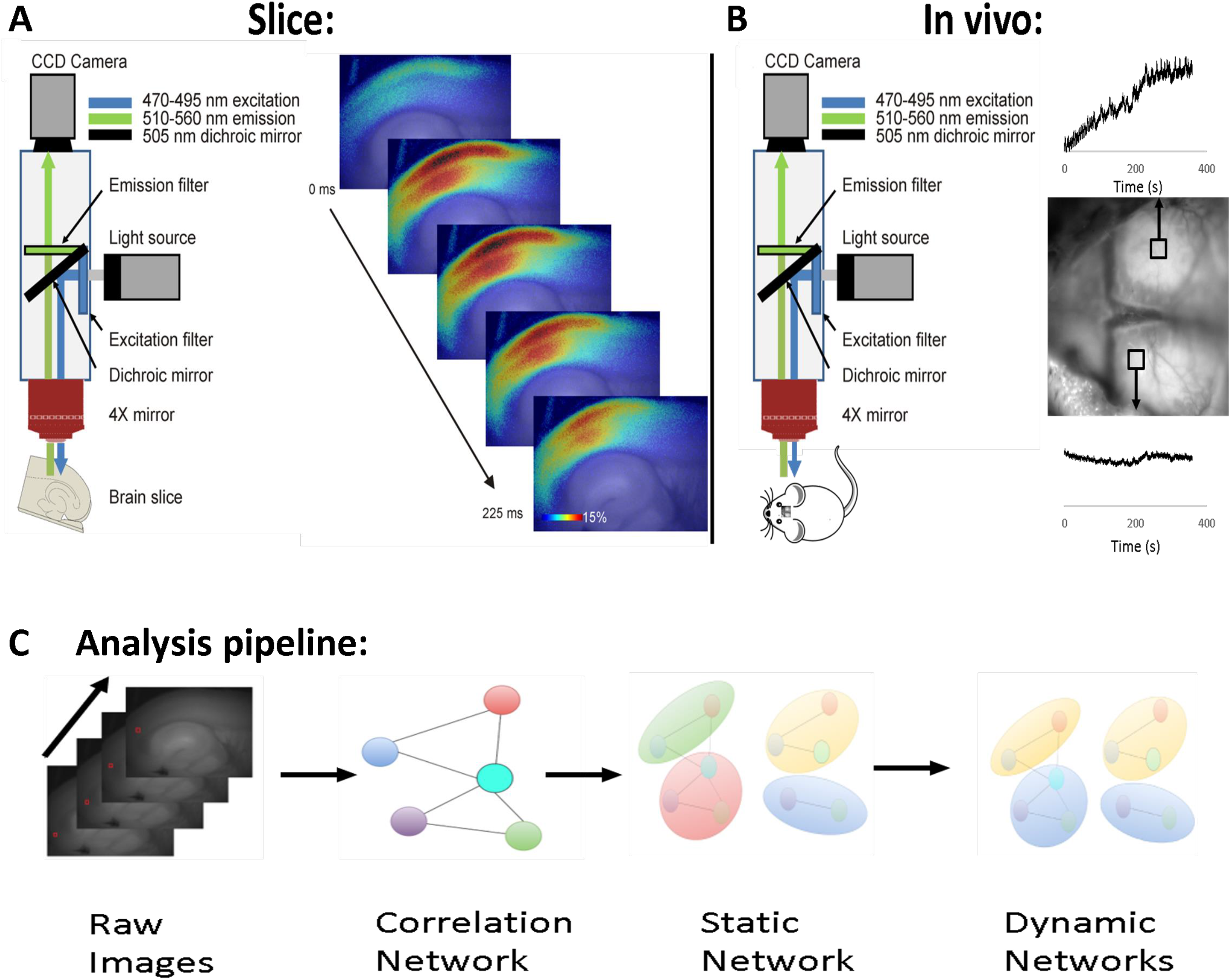
A) Left - Diagram of imaging setup for slice. The brain slice is illuminated with blue light and green light is collected using an epifluorescence setup. Right - A series of images during a paroxysmal depolarization, shown as change in fluorescence over baseline for each pixel. B) Left - Diagram of imaging setup for in vivo experiments. Right – Two representative regions of interest on the surface of the mouse brain, illustrating oscillations in the raw signals. C) Analysis pipeline – 1) Raw images are obtained, 2) Correlation networks are computed, 3) Static networks are determined via Louvain algorithm and 4) CommDy is used to generate dynamic networks.

Prior to applying the CommDy technique, there are several data pre-processing steps required to transform raw brain imaging data into CommDy input, including construction of the time series of a simple network model and static community detection in each time step. After that, CommDy was applied to infer dynamic communities. To create the network model, pixels were used as nodes and a weighted and thresholded network of pixel value correlations across an empirically-determined number of time steps (the “window size”) was then generated. The threshold was determined empirically and correlation values above the threshold were used to build the correlation matrix.

To determine the appropriate time window and correlation thresholds for CommDy analysis applied to flavoprotein autofluorescence imaging data for slice data, the analysis window size was systematically changed from 25 frames (352 msec) to 200 frames (2817 msec) and the resulting spatial distributions of the average degree of each node were examined (See Supplemental Figure 1). For each analysis window (25, 50, 100 or 200 frames), the correlation coefficient for each pair of pixels was computed for the time segment where the analysis window straddled the peak of the response. The average degree of each pixel was then computed as the number of other pixels with which it had correlation coefficients in the range described above each column. Given the large degree of variability seen across images (e.g., there are many more correlations between 0.5-0.6 than 0.9-1.0), the degree of each node was normalized to the maximum degree for each image. Each image, therefore, contains the spatial distribution of normalized node degree for a given analysis window size and given range of correlation coefficient. At an analysis window of 25 frames, high numbers of correlations were seen in portions of the auditory cortex at correlation coefficient thresholds between 0.7 and 0.9, but at thresholds above 0.9, spurious correlations were detected outside of the slice (dashed arrow, top right image). Using a 50 frame window, nodes restricted to the auditory cortex were seen at all thresholds above 0.7. Similar findings were seen at 100 and 200 frames. Given the increased utility of any dynamic network analysis tool at shorter analysis windows (and therefore higher temporal resolution), it was determined that the smallest window not producing spurious correlations would be used for analysis. Similar findings were seen with other activations seen in other animals and a similar analysis was done for *in vivo* data (data not shown). Therefore 50 frames and a correlation threshold of 0.7 (corresponding to the red box in Supplemental Figure 1) were the window and threshold used for all subsequent analyses.

By sliding the correlation window 1 step or frame each time following each correlation extraction, and doing this over the entire timeline, a time series of correlation networks was obtained (Figure 2A-E). CommDy can take the resulting dynamic network as the input directly or, for more efficient processing, it takes any partition of the time steps of the network time series into static communities. In each time step of these time series we applied the Louvain static community inference method [47] to find snapshots of functional clusters, the “groups” that form the basis of CommDy. The Louvain algorithm was chosen because among the modern static community inference methods, its steps are highly intuitive and it is highly scalable, while achieving results comparable to other methods. The algorithm is based on two steps that are repeated iteratively to optimize the modularity in the network ([48], Figure 2F-J). First, the algorithm groups nodes into communities to maximize modularity locally. Then the algorithm hierarchically rebuilds the network aggregating the newly discovered communities into super-nodes. Thus, the nodes in this new network are communities discovered in the previous step and the new links between those nodes are weighted by the cumulative weight of the links between the old nodes that were aggregated into the corresponding communities. The two steps are repeated iteratively until no further increase in modularity is possible.

**Figure 2:**
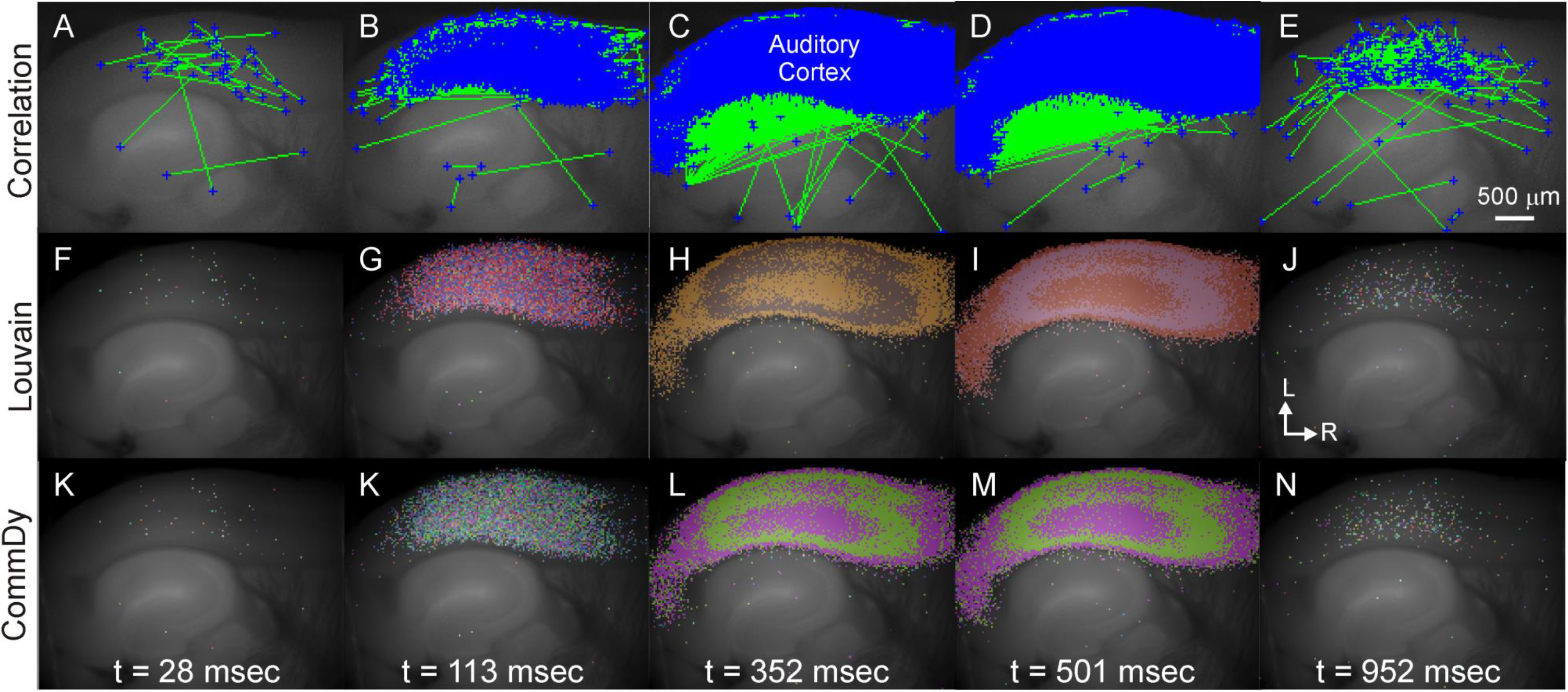
Steps in CommDy, A-E) Sequential images demonstrating all notes (blue symbols) and edges (green line segments), corresponding to all correlations with a correlation coefficient above 0.7.F-J) Networks defined using Louvain algorithm. Each color corresponds to a different network. The networks are reassigned (and recolored) with each frame. K-N) Networks defined using CommDy algorithm. Each color corresponds to a different cluster.

To assess the stability of the Louvain algorithm, which is stochastic in nature, we examined six paroxysmal depolarizations in young mice, and six paroxysmal depolarizations in aged mice. The Louvain algorithm was run 30 times on each paroxysmal depolarization and we computed the standard deviation of each modularity value. We found that all standard deviations were below 5×10^−3^, which is lower than previous reports (Blondel et al. 2008), and, for modularity values above 0.5, the coefficient of variation was below 0.5%, suggesting that these modularity values are stable and the identified static communities are consistent across repetitions. The Louvain algorithm was then compared to the Infomap method [49], which has been used previously to map neural circuits [50, 51]. The algorithm (http://www.mapequation.org/code.html) was run using default parameters, specifying weighted undirected links and optimizing a two-level partition. The input of the algorithm was the correlation network (link list format), while the output is a (.map) file which describes the best two-level partition (no hierarchical structure is preserved). We observed that the Infomap algorithm only detected the large mass of pixels encompassing the auditory cortex, but did not detect heterogeneities in the response seen with Louvain (Supplemental Figure 2).

To understand how the clusters (communities inferred by Louvain algorithm in each time step) change over time, we use the CommDy method [34, 37, 52, 53]. In CommDy, dynamic communities are essentially viewed as dynamic clusters, where the membership of the individual inside the cluster is determined by the total value of the “social cost,” as manifested by the individual’s interactions over time, aiming to minimize interactions outside an individual’s community as well as switching among communities. The inferred community structure parsimoniously minimizes the overall social cost for all the individuals by using the constant-factor approximation algorithm [53]. The definition of social cost is based on two explicit assumptions about individual behavior, motivated by research in social sciences and further supported by the definition of the dynamic community as a cluster. First, it assumes that individuals tend not to change their home community affiliation too often [54]. Second, it assumes that individuals tend to interact with their respective home communities most of the time [55]. These assumptions are translated into three cost parameters potentially incurred by an individual. First, the CommDy method posits a cost for a switch from one community to another. Second, there is a cost of visiting a community of which one is not a member. Third, in data sets for which not all individuals are observed all the time, there is a cost of absence for an individual who is not observed at a gathering of a community of which it is a member. A dynamic community is then defined as a time series of sets of individuals among whom the overall social cost of interacting is minimized [34]. Note that these costs have not yet been defined for brain networks and were set to equal values for our initial studies, with subsequent exploration of the range of the parameters. For visualization, once communities were assigned, the pixels belonging to the top 20 communities over the entire timeline were distinguished using a palette of 20 colors (Figure 2K-N). Spatial relationships between the nodes were not considered when assigning groups because of the potential introduction of bias based on assumptions about the relationship between structural and functional connectivity.

An additional advantage afforded by CommDy compared to other dynamic clustering approaches is CommDy’s ability to quantitatively describe the characteristics of the inferred network activity in terms of the node and structural network metrics based on network analysis theory [36]. These metrics permit quantitative comparisons between different networks (e.g., comparing the cortical networks in aged vs. young animals). Therefore, quantitative analyses of CommDy activation patterns were performed using 10 relevant community metrics that describe the interactions between pixels (see supplementary Table 1 for a listing and definition of the metrics).

### Impact of aging on network dynamics

Paroxysmal depolarizations were imaged in brain slices taken from 5 young (5.5 months, n=5) and 5 aged (22 months, n=5) animals (a total of 41 and 30 activations, respectively). As expected, baseline differences in aged and young animals were noted in hearing thresholds (33.9 ± 2.2 [SD] dB SPL for young and 59.9 ± 27.6 [SD] dB SPL for aged, p=0.032, Mann-Whitney). In addition, as previously described in a related dataset [56], there were differences found in cortical thickness between the two groups (1.20 ± 0.04 [SD] mm for young and 1.04 ± 0.04 [SD] mm for aged, p=0.012, Mann-Whitney).

Initially, static analysis was performed on auditory cortical networks in young and aging mice. To do this, the average degree of each node was computed for each pixel as the average number of edges at each node during the 100 frame window surrounding the peak of the paroxysmal depolarization. These values were then averaged across paroxysmal depolarizations within each animal, normalized to the overall magnitude of the response, and the resulting grand average for each group of animals is shown in Figure 3. As shown, the average normalized degree of each node in all was substantially higher in younger animals than in aged animals (mean normalized node degree in young animals = 813.0 (353.6 SD) connected nodes per active node, mean in aged animals = 149.1 (162.4 SD), connected nodes per active node, p=0.005). These data suggest that the average connectivity between nodes is higher in younger animals than in aging animals, and cannot be accounted for by any differences in the magnitude in the response of the young vs. aged animals.

**Figure 3:**
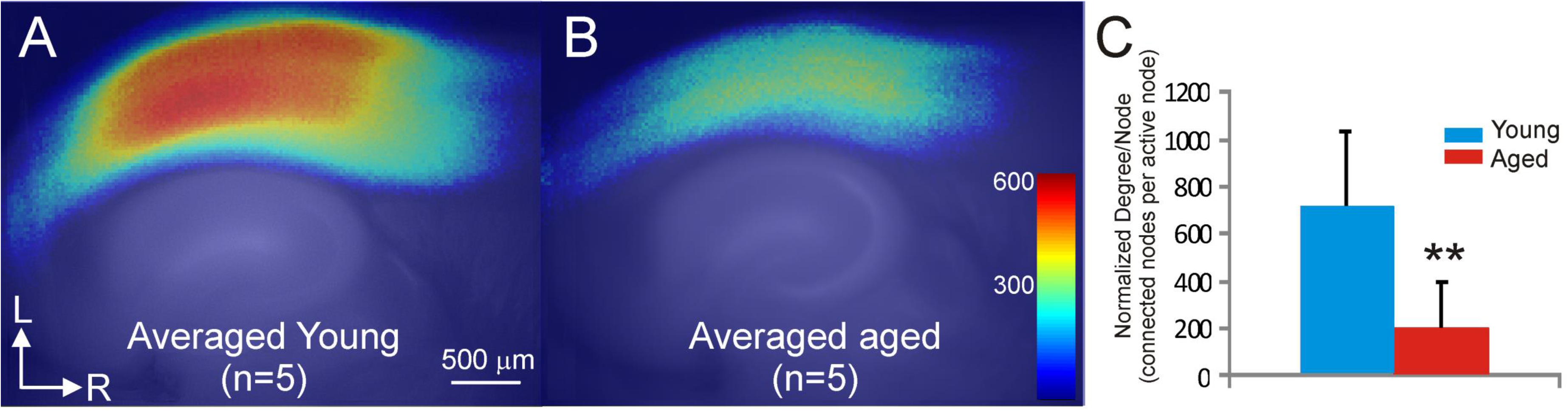
Young animals show greater average degree of each node than older animals. A) Average normalized degree of each node, averaged across all 5 animals, superimposed upon an averaged anatomical map. B) Identical analysis, but for aged animals. C) Bar graph showing that the average degree of each node is significantly higher in the young than the aged animals. Error bars = standard deviation. ** p = 0.005. See text for details.

We then performed dynamic analysis using CommDy, which is built upon static communities detected by the Louvain algorithm independently in each time step. Dynamic analysis of paroxysmal depolarizations in young and aged animals using CommDy revealed that network activity differed between these two groups. Young animals showed distinct patterns of activity across the auditory cortex during paroxysmal depolarizations. See Figures 4A-E for an example from one representative young mouse. Early during the response, multiple small clusters (communities) were observed (Figure 4B). As the response evolves, a distinct cluster (colored red) emerges in the upper layers of the AC (Figure 4C), which then spreads to produce two broad red networks – one in the upper layers and another in the lower layers – which persists to at least 500 msec after onset (Figure 4E, see Supplemental Movie 1). In the aged animals, there is also broad activation across the AC. However, the activation is much more heterogeneous than that seen in the young animal and comprises pixels of many different communities and never coalesces into coherent clusters seen in the young animal (Figures 4F-J for a representative aged mouse, see Supplemental Movie 2).

**Figure 4:**
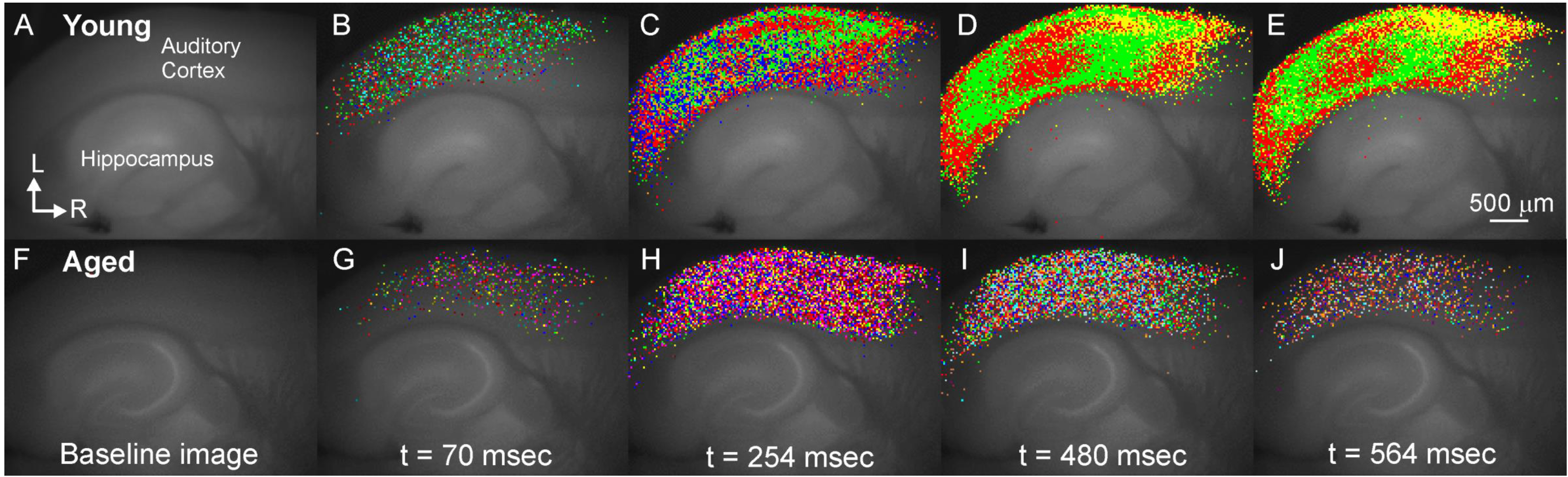
Representative CommDy population dynamics across time for a paroxysmal depolarization in a young animal (A-E) and an aged animal (F-J). Each color represents a different community.

Dynamic network metrics were compared between young and aged mice using univariate and multivariate approaches. To ensure that differences in network measures were not due to changes in the absolute magnitude of activation, individual group and community sizes were normalized by the total number of pixels in each activation. Using a mixed effects model, age was found to be a significant predictor for both normalized group size (mean values = 0.211 ±0.113 [SD] pixels for young vs. 0.058 ± 0.046 [SD] for aged, p = 0.0001) and normalized community size (mean values = 0.235 ± 0.142 [SD] pixels for young vs. 0.046 ± 0.044 [SD] for aged, p = 0.0002). The differences in homing, a measure of network cohesion, trended towards significance but did not survive correction for multiple comparisons (mean values = 0.756 ± 0.663 [SD] for young vs. 0.125 ± 0.081 [SD] for aged, p = 0.0168). See Figure 5A for a comparison of all 10 network parameters.

**Figure 5:**
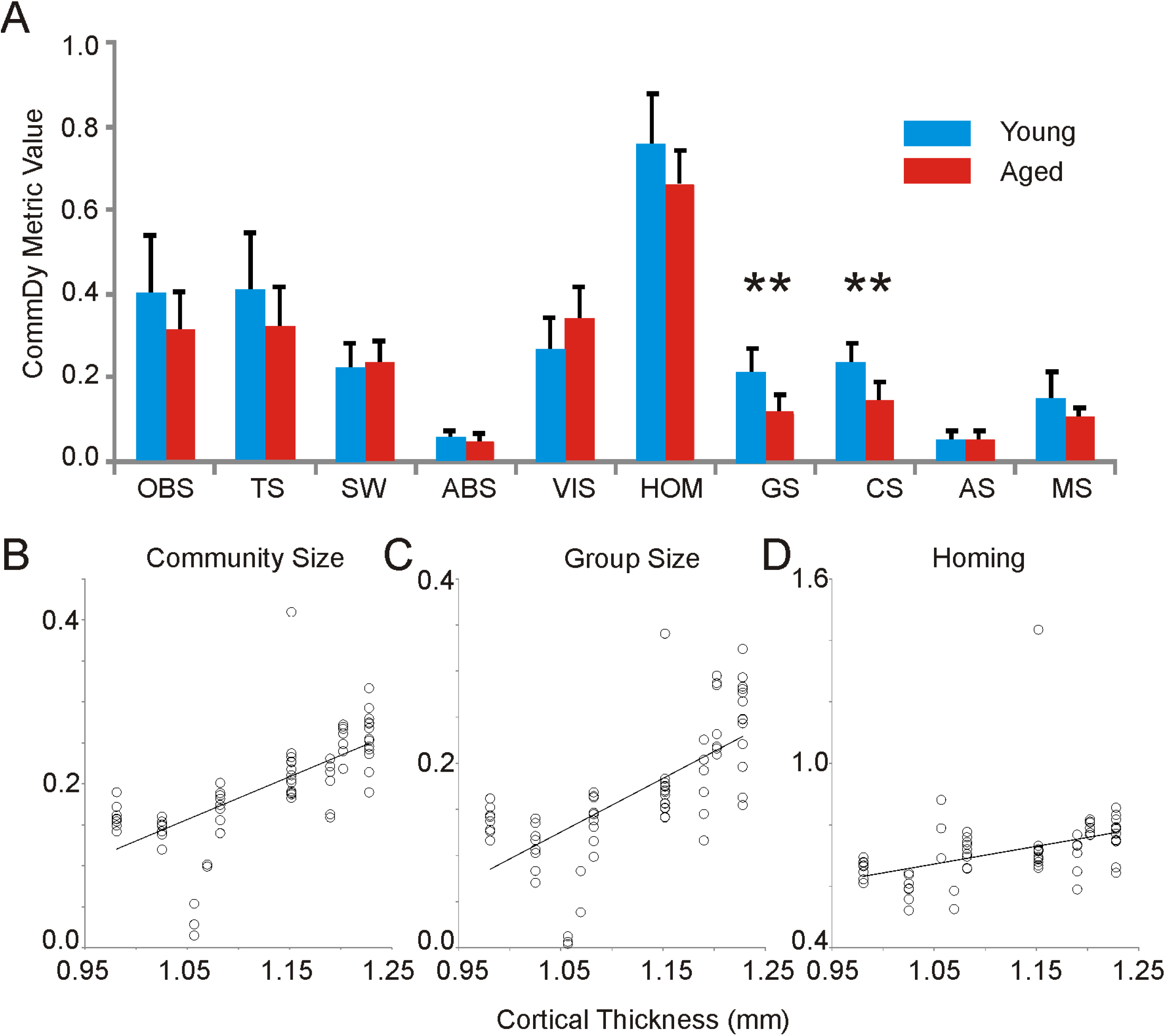
A) Bar graph of average CommDy parameters for young (n=40 activations) and aged (n=31 activations) animals. Group size, community size, average stay and maximum stay are normalized for the total number of activated pixels. Error bars = standard deviation. Correlations between cortical thickness and normalized community size (B), normalized group size (C) and homing (D). The same x-axis is used for all three correlations.

Given the relationship between age, hearing loss and cortical thickness, we examined the relationship between hearing loss and cortical thickness on the various CommDy network parameters. Hearing loss showed no significant relationship to any network metric (lowest p-value = 0.413, observed for maximum community stay), though cortical thickness was significantly correlated to normalized community size (R=0.687, p<0.0001, mixed-effects model, Figure 5B), normalized group size (R=0.687, p<0.0001, Figure 5C), and homing (R=0.423, p=0.0003, Figure 5D).

To examine the relationships among all CommDy variables in young and aged mice, we performed multivariate analysis with PCA, using all activations from all animals. We observed that the first two components account for majority of the variance in the community metrics of all the paroxysmal depolarizations from individual animals. This analysis revealed a clustering of two groups of activations that segregate by age group in the statistical space projected onto the first two components, with little overlap between these groups, shown for each paroxysmal depolarization and coded by individual using different shapes (Figure 6). Analysis of this biplot shows that indicators of network temporal stability and cohesion, such as homing, community stay, group size and community size show strong positive alignment (and are thus correlated) for the cluster of young animals. In contrast, indicators of network fragmentation and instability, such as visiting and switching costs, are more aligned with the cluster for the older individuals. Review of the individual paroxysmal depolarizations from each animal shows that paroxysmal depolarizations from different animals are intermingled, rather than clustered. This finding suggests that the common feature separating the paroxysmal depolarizations is membership in an age group, rather than membership to an individual animal. The weights of all three cost metrics were systematically varied and in all cases, similar separations were seen in young vs. aged animals (data not shown). Taken together, these data suggest that networks from aged animals are more fragmented and less cohesive than those from young animals.

**Figure 6:**
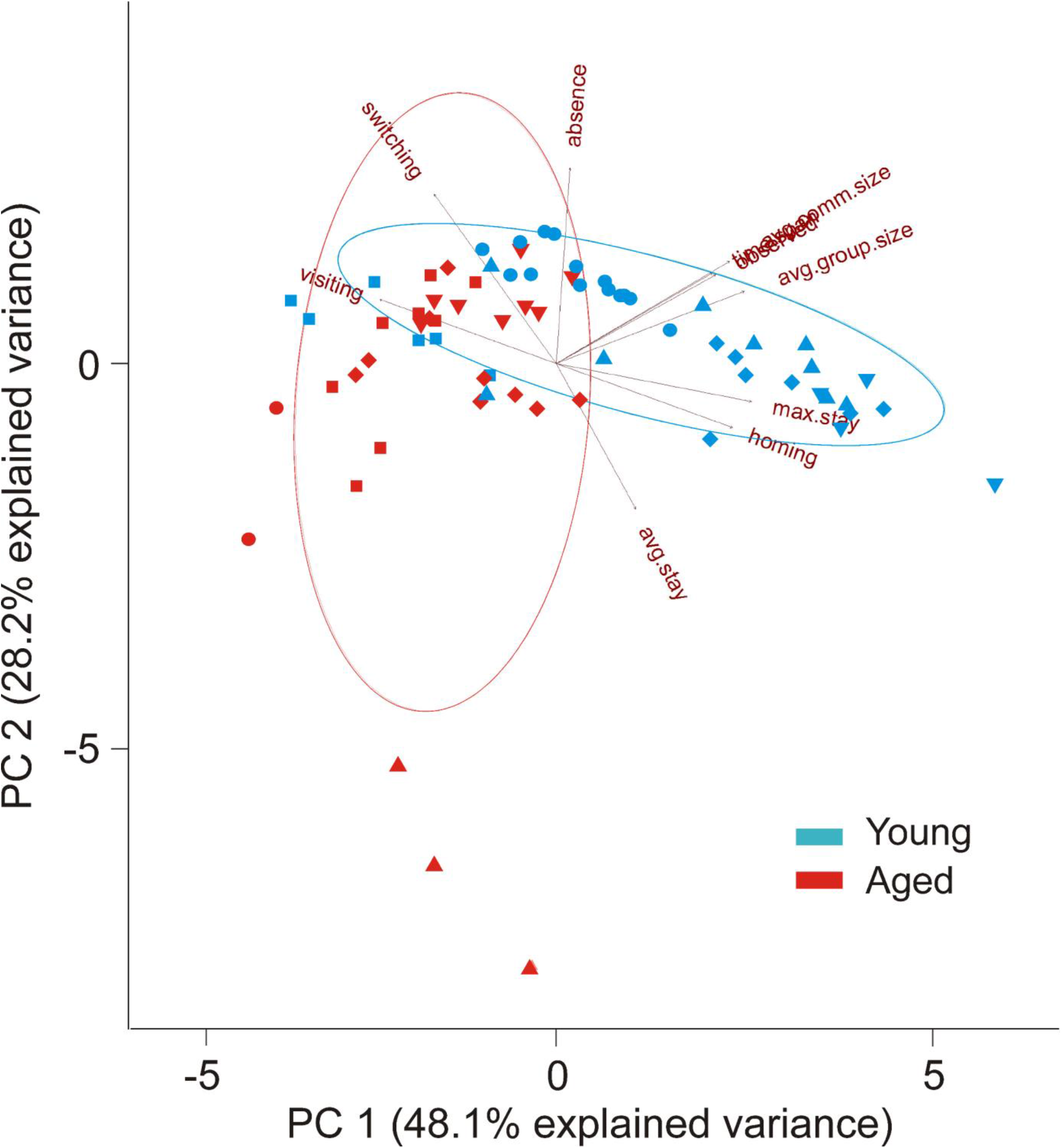
PCA biplot for 10 CommDy parameters for the 71 total activations across 10 animals, each animal coded with a different shape, with old animals in red and young animals in blue. Ellipses correspond to the 95% confidence intervals for each population. Three overlapping parameters in top right are observed, time span and average community size.

To test whether the aging-related changes in group and community size described above may be related to changes in intracortical connectivity, diminished intracortical connectivity was pharmacologically mimicked in a coronal brain slice from a young animal by blocking NMDA receptors, which are enriched in cortico-cortical synapses [57, 58]. A concentration-dependent drop in both normalized group and community size was seen with bath-application of the NMDA blocker APV (1-way ANOVA, p = 5 ×10^−9^ for group size, p = 0.004 for community size, Figures 7A and B). To further determine whether network changes observed during NMDA blockade resembled network changes that occur during aging, a classifier was built using the NMDA blockade data, and the ability of this classifier to distinguish patterns of activity in young vs. aged was measured. A random forest method [59] was used to build a classifier using the five different classes of NMDA blockade data (baseline, 15, 30, 60 and 120 μM APV). Given the 5 dose categories, chance performance of the classifier = 0.2 (or 0.35 using the majority class as the prediction). Using 50 trees and leave-one-out validation, the accuracy of the classifier for APV data = 0.575, which is greater than that expected by chance (see confusion matrix, Figure 7C). When applied to all data obtained from young and aged animals, this classifier identified 30 out of 40 young as belonging to the baseline group and 22 out of 30 aged as belonging to one of the APV groups, thus “correctly” identifying 74% of the activations (Figure 7D). When comparing the 30 μM dose group (the category into which most of the aged data fell) to the baseline group, the overall accuracy of the model was 0.70. Thus, using entirely different datasets for model development and model validation, these data suggest that the aged data, when looked at from a network perspective, bear strong commonalities to the effect of bathing a young slice in 30 μM APV.

**Figure 7:**
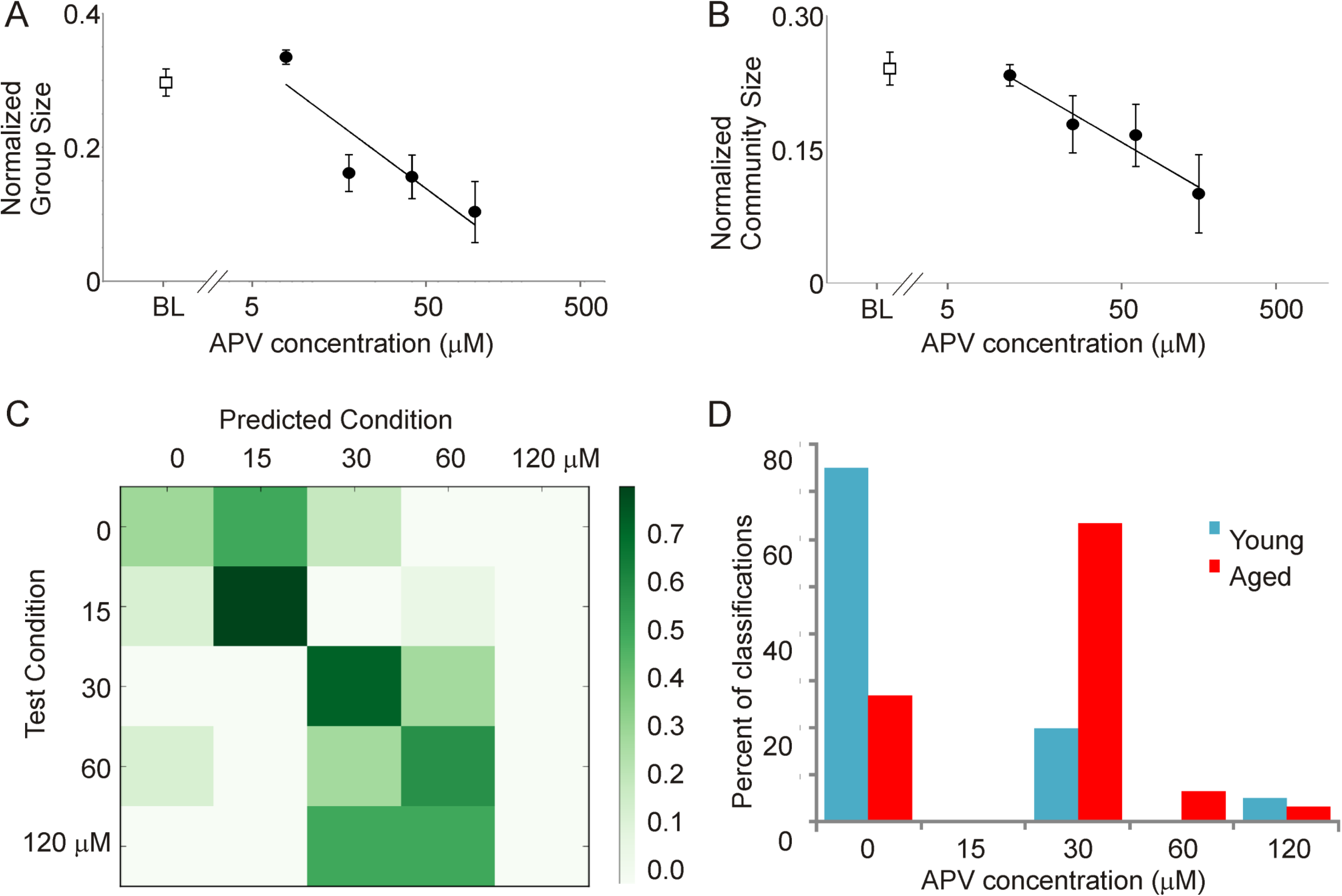
Impact of increasing concentrations of bath-applied APV at concentrations of 15 (n=14 activations), 30 (n=9), 60 (n=9) and 120 μM (n=5) on normalized group size (A) and normalized community size (B) in a coronal slice preparation from a young adult mouse. BL = baseline values prior to adding APV (n = 10). C) Confusion matrix showing the performance of the random forest classifier. True categories are shown in the rows, and predicted category is shown in the columns. Most data are seen on or near the diagonal, yielding an accuracy of 0.575. D) Using the same model to classify data from young vs. aged mice, most activations from young animals (30/40) fall correctly into the baseline (no drug) category, while most of the activations from aged animals (22/30) fall into a drug category.

### Analysis of in vivo data

9 young (average age = 4.7 months, range = 4.4 to 5.0 months) and 9 aged (average age = 23.2 months, range = 23.0 to 23.5 months) mice were used for these experiments. Since flavoprotein autofluorescence has not been used previously to measure spontaneous neural activity *in vivo*, we initially characterized the spectral properties of the signals. The majority of the power is in the delta range (approximately 3 Hz) in both young and aged animals. To determine the modulation of spontaneous activity by anesthesia, mice were anesthetized with ketamine/xylazine (after spontaneous signals were measured for CommDy analysis) and we observed the expected overall decrease in frequency in both groups (p=0.006), as has been seen previously [60, 61], with no differences between groups (p=0.3, Supplemental Figure 2B). These data suggest that spontaneous flavoprotein autofluorescence signals track with more commonly-measured indicators of spontaneous activity, such as EEG, but given the slow nature of the signal are limited to lower frequency spectral bands than EEG.

To compare the overall degree of connectedness between nodes in aged vs. young animals, the degree of each node was computed. Similar to the findings in brain slices from the auditory cortex, the degree of connectivity, as measured by average number of edges per node, was significantly decreased in the aged compared to the young animals (156.7 × 10^4^(SE 53.2 × 10^4^) edges/node vs. 6.1 × 10^4^(SE 3.8 × 10^4^) edges/node, p = 0.02, See Figures 8A and B).

**Figure 8:**
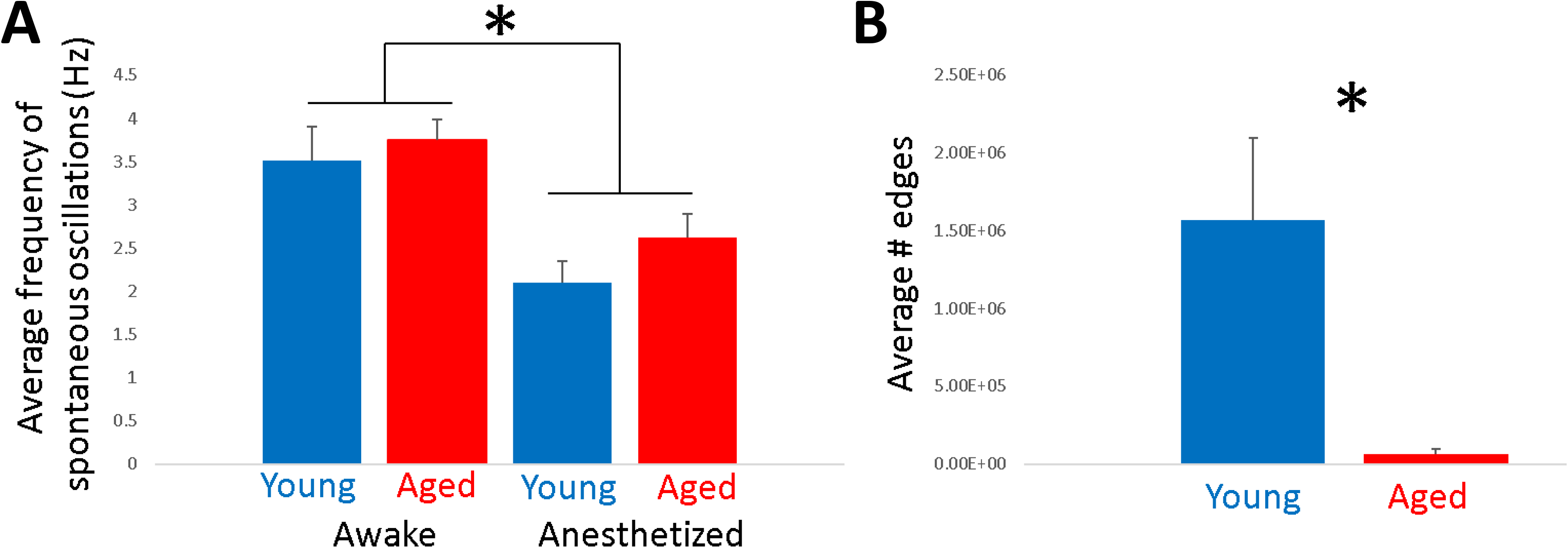
A) The average frequency of spontaneous activity was roughly 3.5 Hz in awake mice and similar between aged and young mice. The average frequency declined significantly to approximately 2.5 Hz in the presence of ketamine/xylazine anesthesia. B) Despite the similarity in oscillatory activity, aged animals had a marked decrease in cortical connectivity, as measured by the average number of edges in the correlation networks.

## Discussion

The notion that neuronal cell assemblies are critical for perception and may shift dynamically over time is a foundational principle in neuroscience [62, 63], but quantifying such network dynamics has been challenging. Here we provide proof-of-concept data that dynamic network analysis, specifically CommDy, a network analytical technique initially developed for the inference of communities in dynamic social networks, can quantitatively describe network-level dynamics in brain imaging experiments. It is further shown that CommDy analysis from two different brain regions under different conditions (slice vs. *in vivo*) shows that aging is associated with a series of changes in network metrics that demonstrate aging-dependent fragmentation of neuronal activity. It is further shown that the network characteristics seen are similar to those seen after partial NMDA receptor blockade, suggesting a possible mechanism for this aging-related fragmentation. Below we describe the limitations and the potential utilities of the CommDy technique in brain studies and how they relate to changes occurring during aging.

### Technical considerations

The current work is based on flavoprotein autofluorescence imaging, which has the advantages of providing a stable intrinsic signal with high sensitivity and good spatial resolution (100-200 microns [38]), but has a relatively sluggish time course (~0.5-1 sec for time to peak [38–40]). Given this slow time course, one might expect that small differences in the time to activate of different groups of neurons would be washed out by delays induced by neurometabolic coupling. However, despite these slow responses, CommDy did reveal distinct populations, suggesting that more subtle heterogeneities than could be appreciated using traditional analytical approaches were detectable using CommDy in the time courses of the imaging responses.

Another potential concern about the current analysis is the use of pixels as individual nodes, rather than using defined regions or individual neurons. At the binning and magnification used in this study, an individual pixel occupies an area of approximately 800-1600 μm^2^. To estimate the number of neurons represented in this pixel, an upper estimate of the volume will be used, corresponding to the full thickness of the slice or the cortex, though it is likely that the blue excitation light incompletely penetrated the sample. The mouse cerebral cortex contains about 9.2×10^4^ neurons/mm^3^ [64] giving a rough estimate that each pixel in our preparation contained signal emanating from on the order of ~50-100 neurons. Compared to techniques that image spiking activity of individual cells, such as calcium imaging, flavoprotein autofluorescence offers lower spatial resolution but offers a broad field of view (multiple millimeters), permitting large-scale networks to be identified. Compared to the most-commonly used technique for large-scale brain imaging, fMRI with BOLD signals, where the voxel sizes range from millimeters in humans to hundreds of microns in rats, the current analysis provides substantially higher spatial resolution. It is important to note that the current analytical approach may be easily adapted to any of the imaging techniques mentioned above.

### Implications for aging

Pathological changes associated with aging have been described at virtually every level of the central nervous system, including the cortex. Here, we observed aging-related disruptions in network-level function of the auditory and motor cortices. In the slice preparation, these changes are independent of the magnitude of the activation, and are likely not related to differences in slice viability, since we have recently found that the auditory cortices in slices from young vs. aged animals do not differ in their baseline redox state or response to metabolic stress [56], and are also likely not related to differences in GABAergic synaptic inhibition, since saturating doses of a GABA_A_ antagonist were used in the slice study. Similar fragmentation of networks was also observed in the motor cortex, despite preservation of the overall spectral structure and spectral responsiveness of the flavoprotein signal. Therefore, the current data suggest that aging causes changes in the underlying local excitatory substructure in the cerebral cortex. The specific substrate for these changes is unknown, but could relate to changes in local dendritic branching and integration, changes in intrinsic excitability of pyramidal cells, or changes in synaptic properties, all known to be altered in the aging brain [65–68]. Regarding the latter hypothesis, we observed that a classifier built only using dose-dependent changes in CommDy network parameters induced by APV was able to classify data from young vs. aged animals with a high degree of accuracy (Figure 8). These findings are remarkable in that the cross-validation was done using entirely different datasets obtained from different types of slices (auditory cortex slices for aging data and coronal slice for APV data). Further, these findings are consistent with a body of literature demonstrating NMDA receptor hypofunction in the cerebral cortex of the aging brain [69–71]. Therefore, these data suggest that the aging-related changes in network-level interactions observed in the cortex of this and previous studies [20, 21] may be, at least in part, caused by changes in NMDA receptor function.

Another novel finding in this study is the observation that cortical thickness is a significant predictor of group size, community size and homing in the auditory cortex (Figures 5B-D). This relationship is consistent with the mechanisms of aging-associated thinning of the cortex. Cortical thinning is related to the loss of synapses between cortical neurons and/or demyelination of local cortical axons, with retention of the number of neurons [10, 72–74]. As such, cortical thinning would be predicted to be associated with drops in cortical connectivity, consistent with the current data. This idea is supported by previous findings from network theory showing that random or uniform loss of connectivity, translated into uniformly lower network density, results in smaller clusters [75, 76]. Hence, our results showing diminished group and community size with aging, combined with the known drop in synaptic connectivity seen in aging, are consistent with predictions from network theory.

In addition to aging-related decreases in network size, multivariate analysis revealed aging-related changes in virtually all metrics of network cohesion (e.g., visiting, switching, homing, maximum community stay, Figure 5). In this context, it is worth pointing out that simple loss of connectivity does not by itself directly imply lack of cohesion over time, as the small groups can still coalesce into functional units. For example, even at low densities, the presence of well-connected clusters has been shown to result in synchronization in static networks [77–79]. Thus, our results that show lower cohesion measures in older brains indicate that the loss of cohesiveness cannot only be explained by drops in network connectivity and requires further investigation.

### Potential utilities of CommDy

CommDy may prove to be a useful tool to quantitatively study brain networks. Although the current study is focused on flavoprotein autofluorescence imaging, CommDy can be adapted to other forms of brain imaging as well. Such adaptation would permit CommDy to be used to refine network-level hypotheses about brain function, for example, in the study of pathological states in humans. It is speculated that many neurological and psychiatric disorders are caused by functional disruptions in large-scale brain networks [80–82]. Most of the current work in this area has focused on using network measures either as a tool to classify different disease states, or as a means to better understand network disruptions associated with disease states. In both cases, much of the work has been focused on the generation of static maps. However, in brain diseases, fluctuations in clinical symptomatology are the rule rather than the exception. For example, seizures in epilepsy, “off” states in Parkinson’s Disease and hallucinations in schizophrenia, all occur paroxysmally on the backdrop of stable brain structures, such that it will only be possible to understand them using dynamic assessment tools. CommDy may add to the growing toolbox of network assessment tools and will contribute to the understanding of brain network dynamics.

## Methods

### Animal use

CBA/CaJ mice from 4.4 to 23.5 months of age of both sexes were used. All procedures were approved by the Institutional Animal Care and Use Committee at the University of Illinois. All animals were housed in animal care facilities approved by the American Association for Assessment and Accreditation of Laboratory Animal Care.

### Auditory brainstem responses (ABRs)

As previously described [56], to measure hearing for the aging studies, ABRs were obtained in response to tones at frequencies of 4, 8, 16, 32, 45, and 64 kHz, as well as in response to broadband noise. Animals were anesthetized with 100 mg/kg ketamine + 3 mg/kg xylazine intraperitoneally before the insertion of two subdermal electrodes, one at the vertex and one behind the left ear. Stimuli were presented using a Tucker-Davis (TDT) system 3, ES1 free field speaker, with waveforms being generated by RPvdsEx software. The output of the TDT speaker was calibrated at all the relevant frequencies, using a Bruel and Kjaer type 4135 microphone and a Bruel and Kjaer measuring amplifier (Model 2610). Each frequency was presented for 5ms (3ms flat with 1ms for both rise and fall times), at a rate of 2-6Hz with a 100ms analysis window. Raw potentials were obtained with a Dagan 2400A amplifier and preamplifier headstage combination, and filtered between 100Hz and 3000Hz. An AD instruments PowerLab 4/30 system was used to average these waveforms 500 times. Significant deflections, assessed via visual inspection, within 10ms after the end of the stimulus were deemed to be a response. Blinding of ABR assessments was not done given the conspicuous physical differences in young vs. aged mice (i.e., aged animals are heavier and had less hair on snout).

### Brain slice preparation

The CBA/CaJ strain was used because of its gradual aging-related hearing loss [83–85] Five young mice (5.5 months of age) and five aged mice (22 months of age) were used. Mice were initially anesthetized with ketamine (100 mg/kg) and xylazine (3 mg/kg) and then transcardially perfused with an ice-cold sucrose saline solution (in mM: 206 sucrose, 10.0 MgCl_2_, 11.0 glucose, 1.25 NaH_2_PO_4_, 26 NaHCO_3_, 0.5 CaCl_2_, 2.5 KCl, pH 7.4). Slices containing the auditory cortex were cut using a modification of the method developed by Cruikshank et al. [86], modified for the aging brain, as described previously [56, 87], see Figure 1B for brain image. Brains were blocked by removing of the olfactory bulbs and the anterior 2 mm of frontal cortex with a razor blade. The brain was then tipped onto the coronal cut and an off-horizontal cut was made on the dorsal surface, removing a sliver of brain angled at 20° from the horizontal plane. The brain was then glued onto the cut angled surface, and sections were then taken. All slices were then transferred to a holding chamber containing oxygenated incubation artificial cerebrospinal fluid (ACSF) (in mM: 126 NaCl, 3.0 MgCl_2_, 10.0 glucose, 1.25 NaH_2_PO_4_, 26 NaHCO_3_, 1.0 CaCl_2_, 2.5 KCl, pH 7.4) and incubated at 32°C for 1h prior to experimentation. The ACSF used for the experiments use equimolar MgCl_2_ and CaCl_2_ and contained 1 μM of the GABA_A_ antagonist SR95531 to enhance spontaneous activity. Without the addition of SR95531, we found that the amount of activity present in the slice was insufficient for either CommDy or traditional analyses. During imaging, slices were placed on a stainless steel mesh for 2-sided perfusion, as we have described previously [88]. To assess the impact of NMDA blockade on network activity, the NMDA blocker D-APV (Tocris, catalog# 0106) was dissolved in ACSF and bath applied at a series of concentrations ranging from 15 to 120 μM, with 20 minute wash-ins between each dose escalation. Cortical thickness was evaluated by drawing a line tangent to the rostral-most extent of the hippocampus, from the white/grey matter border of the cortex to the pia, as we have described previously [56].

### In vivo preparation

An initial stereotactic surgery under ketamine/xylazine (100 mg/kg and 3 mg/kg IP, respectively) was done to glue a threaded headbolt onto the skull using OptiBond XTR kit (Cat # 35106) cement. After at least 3 days to recover, mice were gradually acclimated to an imaging chamber within a soundproof booth. Their headbolts were affixed to a holder, and the body was suspended while under isoflurane (4%) anesthesia. The mice were then allowed to emerge from anesthesia, initially for 5 min, and gradually up to 15-20 min. Awake data were obtained once isoflurane was removed for at least 10 minutes. Once awake data were obtained, the animal was re-anesthetized with ketamine/xylazine (100 mg/kg and 3 mg/kg IP, respectively) for additional imaging.

### Imaging

For the slice work, flavoprotein autofluorescence imaging was done with a fluorescence illuminator (Prior Lumen 200) and a UMNIB Olympus filter cube (470–490 nm excitation, 505 nm dichroic, 515 nm emission long pass). A coverslip was placed over the slice to provide a stable imaging plane and data were collected using an infinity-corrected 4X macro objective (NA 0.28) and a Retiga EXi camera and StreamPix software (See Figure 1A for diagram of experimental setup). Data were obtained with 8×8 hardware binning, producing images of 130 × 174 pixels. Acquisition rates were 71 frames per second.

*In vivo* imaging was done using a macroscope system outfitted with 85 mm f/1.4 and f/1.2 Nikon lenses and an Adimec 1000m CCD camera (7.4 × 7.4 μm pixel size, 1004 × 1004 pixels). Blue light (450 nm, 30 nm band-pass) was used for excitation and green light (515 nm, long-pass) was collected, and a 495 DRLP dichroic mirror was used. Images were collected at 25 frames per second. Pixels were binned 4×4 and a region of interest containing 60 × 70 pixels over both motor cortices and symmetric with respect to the midline was selected for analysis. No filtering was done on the signals.

### Paroxysmal depolarizations

In the presence of SR95531, spontaneous activations were observed with flavoprotein autofluorescence imaging, consisting of a relatively sharp rise in fluorescence, peaking at about 150 msec after baseline, with a slow decline, gradually returning to baseline seconds (Figure 1A, red trace). We routinely observe these spontaneous activations in all brain slices that contain the cerebral cortex and are bathed in concentrations of SR95531 above 0.5 μM and generally such activations occur roughly every 30-60 seconds with no obvious periodicity. These electrical events strongly resemble the intra- and extracellular manifestations of paroxysmal depolarizing shifts, respectively [89–92], and we have previously found that they correlate with paroxysmal depolarizing shifts [93]. Therefore, we refer to them as paroxysmal depolarizations. All slice analyses in this report are done on activity occurring during the paroxysmal depolarizations.

### Statistical analysis

To examine the relationships between 10 CommDy network metrics described in Table 1 across different pixels or across different groups, principal components analysis (PCA) was used. PCA was run in R [94]. To compare activations in slices from young vs. aged mice, a linear mixed models analysis was run in SAS, incorporating random effects because multiple paroxysmal depolarizations occurred in some slices. The 10 CommDy network parameters were used as regression variables and young vs. aged was used as treatment groups. The Holm-Bonferroni method [95] was used to adjust for multiple comparisons.

**Table 1:**
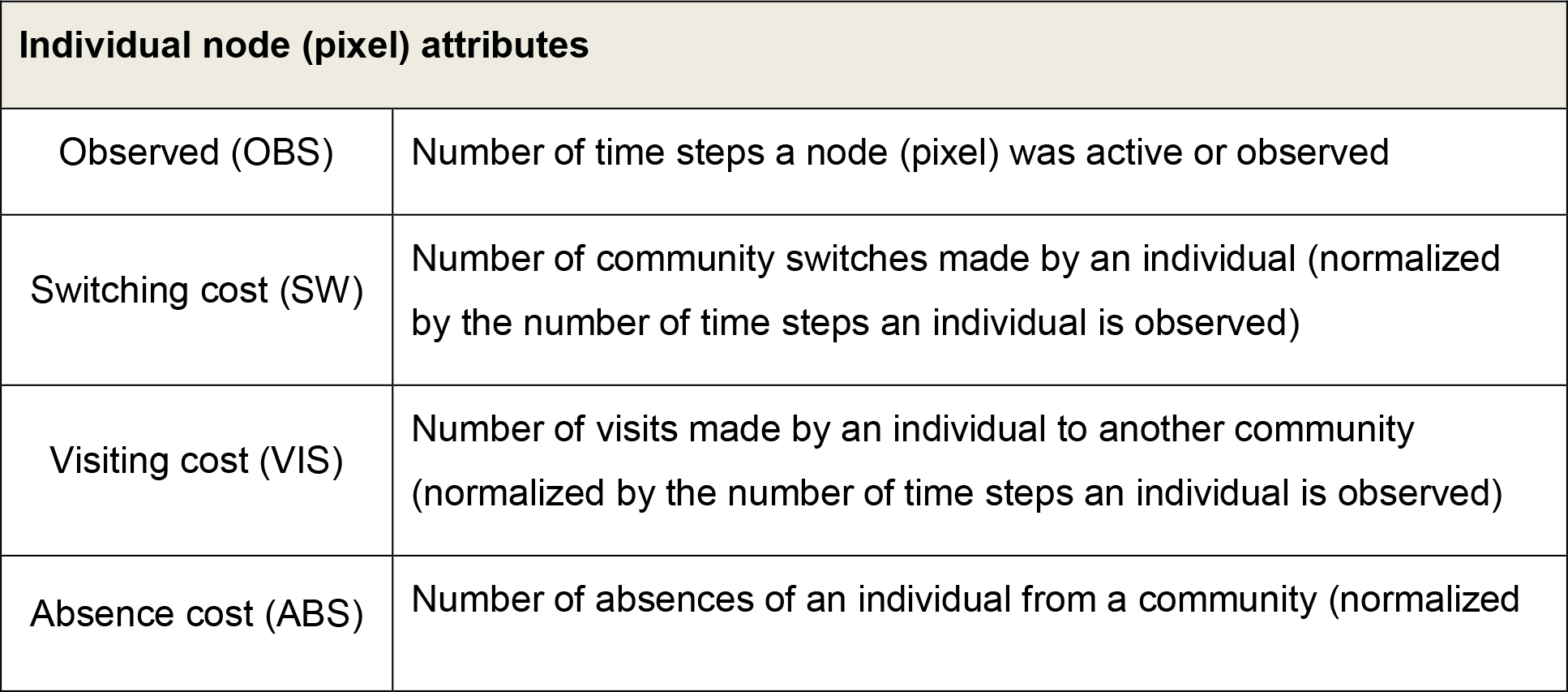
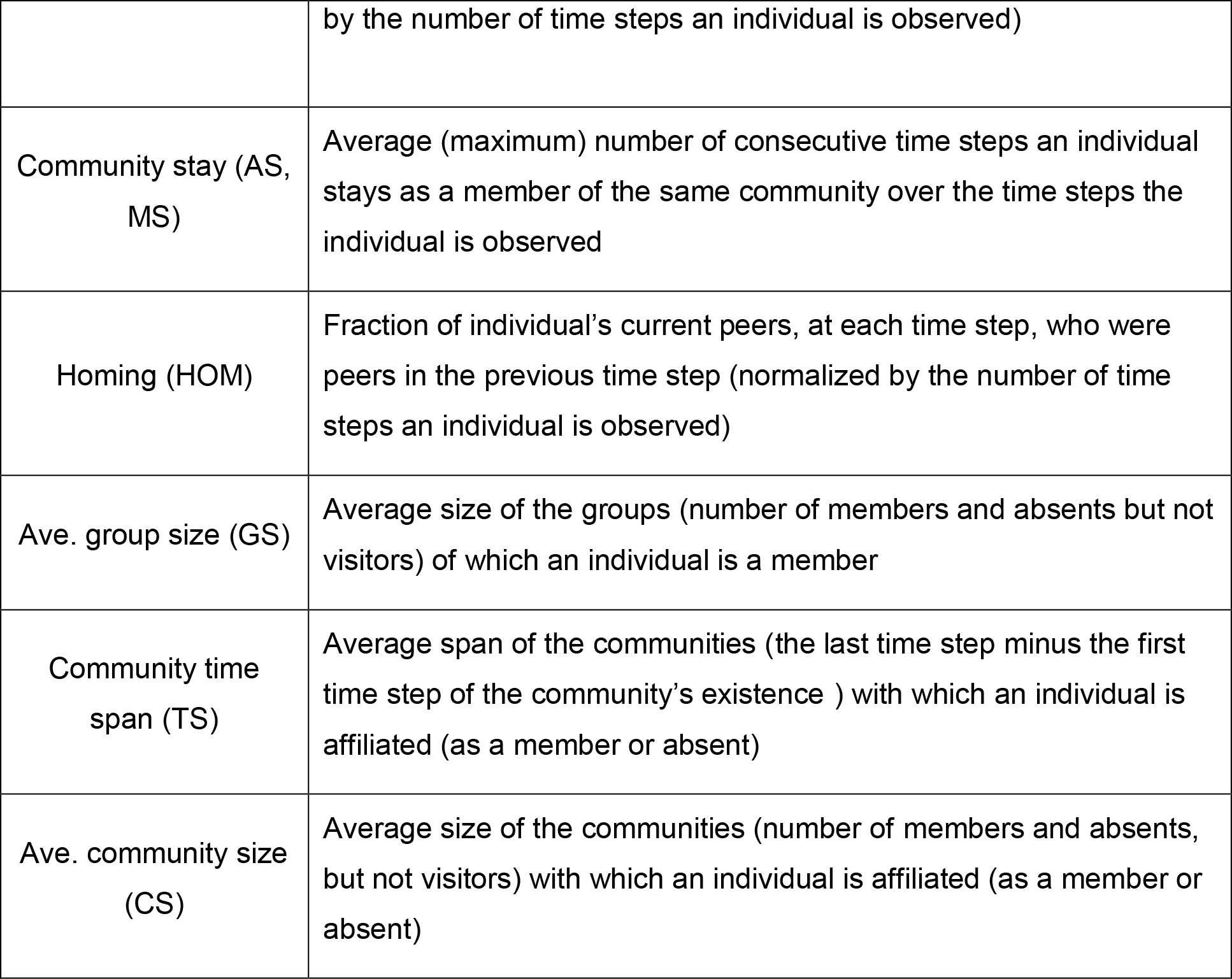
Descriptions of network metrics

## Acknowledgments

This work was supported by NIH DC012125, AG059103, NSF CRCNS grant 1515587, the American Federation of Aging Research and the Alzheimer Association. The authors thank David Trinco and Syed Haider for their data analysis.

---

**Supplemental Figure 1:**
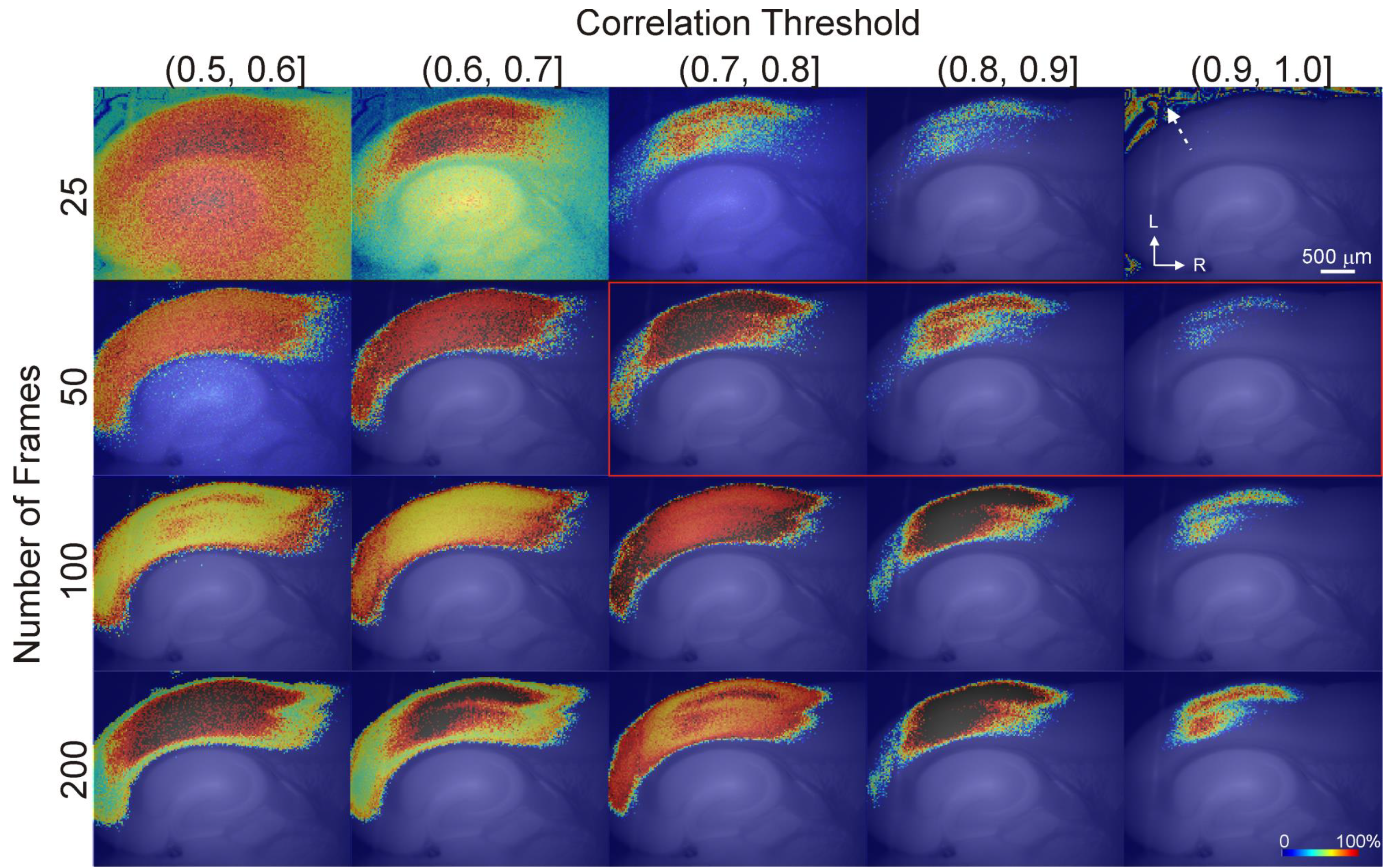
Measurement of the temporal resolution of CommDy was done by varying the length of the sliding window used for analysis, and assessing the resulting spatial distribution of the degree of each node. Each image shows the normalized degree of each node, computed as the number of other pixels with which each node shares a correlation coefficient value between values shown at the top of each column. The degree of each node was computed for different temporal windows, shown at the left of each row. All node degree values were normalized and are expressed between 0-100% (colorbar). Intervals of correlation coefficients are exclusive of the lower number and inclusive of the higher number. The dotted arrow shows an area of spurious correlations outside of the brain, seen with the shortest window length. Red box corresponds to the frame rate and correlation coefficients used in this study.

**Supplemental Figure 2:**
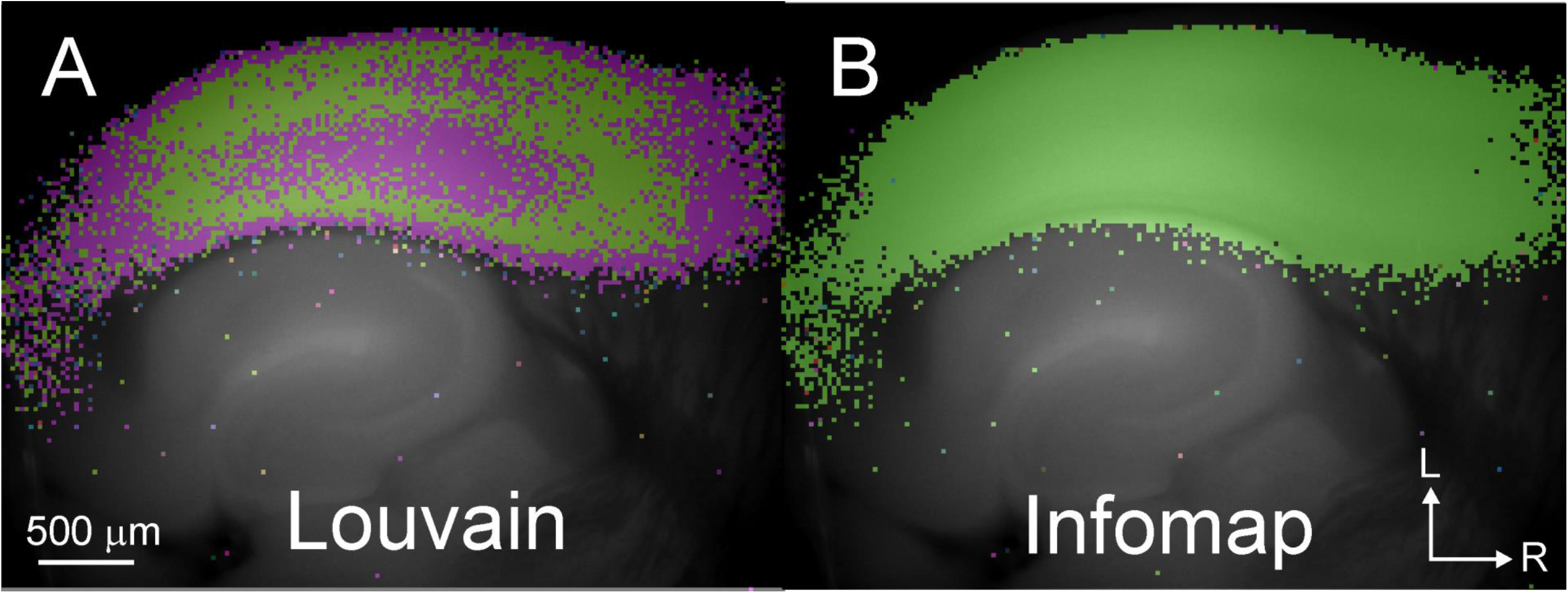
Louvain algorithm is better able to detect subnetworks in this dataset than Infomap. Left – Networks identified in a young animal by the Louvain algorithm near the midpoint of the paroxysmal depolarization. Right – Analysis of the same data using the Infomap algorithm at the same timepoint.

